# Xenon LFP Analysis Platform is a Novel Graphical User Interface for Analysis of Local Field Potential from Large-Scale MEA Recordings

**DOI:** 10.1101/2022.03.25.485521

**Authors:** Arjun Mahadevan, Neela K. Codadu, R. Ryley Parrish

## Abstract

High-density multi-electrode array (HD-MEA) has enabled neuronal measurements at high spatial resolution to record local field potentials (LFP), extracellular spikes, and network activity with ease. Whilst we have advanced recording systems with over 4000 electrodes, capable of recording data at over 20 kHz, it still presents computational challenges to handle, process, extract, and view information from these large recordings. It can be challenging for researchers to extract and view even a single channel that has more than a billion data points, let alone process a group of channels.

We have created a computational method, and an open-source toolkit built on Python, rendered on a web browser using Plotly’s Dash for extracting and viewing the data, and creating interactive visualization. In addition to extracting and viewing entire or small chunks of data sampled at lower or higher frequencies, respectively, it provides a framework to collect user inputs, analyze channel groups, generate raster plots, view quick summary measures for LFP activity, detect and isolate noise channels, and generate plots and visualization in both time and frequency domain. Incorporated into our Graphical User Interface (GUI), we also have created a novel seizure detection method, which can be used to detect the onset of seizures in all or a selected group of channels and provide the following measures of seizures: distance, duration, and propagation across the region of interest.

We demonstrate the utility of this toolkit, using datasets collected from the 3Brain BioCAM duplex system. For the current analysis, we demonstrate the toolkit and methods with a low sampling frequency dataset (300 Hz) and a group of approximately 400 channels. Using this toolkit, we present novel data demonstrating increased seizure propagation speed from slices of Scn1aHet mice compared to littermate controls.

With advances in HD-MEA recording systems with high spatial and temporal resolution, limited tools are available for researchers to view and process these big datasets. We now provide a user-friendly toolkit to analyze LFP activity obtained from large-scale MEA recordings with translatable applications to EEG recordings, and demonstrate the utility of this new graphic user interface with novel biological findings.

## 1. Introduction

The technology of neuronal data acquisition using high density multi-electrode arrays (HD-MEAs) in tissue and cell cultures has grown dramatically over the past decade (Maccione et al., 2013, Maccione et al., 2015, Ingebrandt, 2015, Müller et al., 2015, Steinmetz et al., 2019, Paulk et al., 2022, Maccione et al., 2014). These ever-growing state-of-the-art electrophysiology techniques (Miccoli et al., 2019, Lopez et al., 2018, Stevenson and Kording, 2011) now include commercially available HD-MEA devices that can record extracellular neuronal signals from cell cultures or brain slices on 6 wells simultaneously, where each well consists of 1024 electrodes for a total of 6144 electrodes sampled at 20 kHz or 2304 electrodes for a total of 13824 electrodes at 10 kHz (3BrainAG, 2022). A single recording can range in file size from 5GB/minute or larger compressed. Several pharmaceutical applications and drug-testing protocols require long-duration recordings from 45 minutes to 90 minutes (Codadu et al., 2019a), which can result in large data files of 350 to 500 GB from these recordings.

While electrophysiology and chip technology has progressed at a rapid pace generating high-quality precise neuronal data with a high degree of spatial accuracy, developing data analysis platforms and algorithms exploiting the full potential of the recordings is quite challenging (Mahmud and Vassanelli, 2016, Paninski and Cunningham, 2018). The progress in data analysis pipelines, big data algorithms, flexible analysis platforms to adapt to different techniques, data formats, and research requirements is slowly evolving to handle the large scale of data (Landhuis, 2017). Most applications using high density MEA recordings rely on analysis of high-frequency activity, such as action-potential data, to include useful features, such as spike sorting, clustering, and classification, which has received a lot of attention in the research community, including several open-source architecture toolboxes to view and process the data (Pachitariu et al., 2016, Yger et al., 2018, Lee et al., 2020). Proprietary software and open-source toolboxes that come with the HD-MEA measurement systems can sometimes be restrictive to researchers. While they do provide blackbox-type solutions to spike identification, sorting, generating raster plots and other measures, they may not offer enough customization and adaptability to different methods of viewing and analyzing the data (Bridges et al., 2018). Moreover, while different toolboxes and software platforms provide different functionality, there are benefits and limitations related to the scalability of algorithms for large-scale data, and new paradigms are constantly evolving to exploit the vast potential of these recordings (Mahmud et al., 2012, Diggelmann et al., 2018, Sedaghat-Nejad et al., 2021, Buccino, 2022 January 7).

There appears to be many options for analysis of high-frequency activity for large-scale MEA recordings (Peter C. Petersen, 2021, Xin Hu, 2022, Franke et al., 2015, Buccino et al., 2020). However, open-source, user-friendly analysis platforms for visualizing long recordings of LFP collected from HD-MEA systems is limited. From our review of literature and open-source toolboxes, there are limited data-analysis pipelines that are flexible, customizable, and object-oriented methods for processing and visualizing data for low-frequency (0.5 to 300 Hz) LFP activity. This will continue to limit the usefulness of these large-scale MEA recording systems for many electrophysiologists. Nevertheless, there is an increasing number of research labs using HD-MEAs to record LFP activity to understand neuronal network dynamics from cortical slices (Hu et al., 2022, Medrihan et al., 2015, Toader et al., 2013, Ferrea et al., 2012). One available toolbox to view and process MEA data is presented by Bridges et al. (Bridges et al., 2018), built using Python leveraging GPU (Graphics Processing Units) capabilities to view and generate visualization for large MEA data files. In our current work, we present a much different data pipeline with diverse features and summary metrics, that is built on Python, rendered on a browser using Plotly’s Dash. This data-analysis pipeline is for band-pass filtered (0.5 to 2048 Hz) LFP activity and seizure analysis that is scalable to large datasets, with an interactive GUI for analyzing HD MEA measurements. This GUI includes several features to generate summary measures and plots, and trace LFP activity over time. For people familiar with basic Python, this can also serve as a boiler plate to customize, and add functions and visualization based on individual researchers’ analysis requirements.

Researchers also require novel ways to track LFP activity over space and time, as calcium imaging is limited by slow kinetics (Wei et al., 2020, Vanwalleghem et al., 2020, Helassa et al., 2016, Tang et al., 2015) and current voltage-imaging techniques have several weaknesses, such as high-bleaching properties (Xiao et al., 2021, Kulkarni and Miller, 2017). Recordings using high-resolution MEA systems offer a new way to explore network communication, with high degree of time and spatial resolution, but require tools to tap into their full potential. Our new data pipeline offers an efficient and easy tool to analyze the spatial and time resolution offered by these MEA systems. We demonstrate the utility of this data pipeline with induction of seizure-like activity and generating example LFP raster plots over time and space, along with example traces from subregions of the brain. This bird’s-eye view of LFP activity within our GUI creates a new tool for investigation into novel insight into network dynamics such as how neocortex and hippocampus interact with each other. Furthermore, we demonstrate a novel seizure-tracking approach using the high density of electrophysiological channels with potential to be superior to large-scale calcium imaging to track seizure dynamics. We present data using this analysis tool that shows brain slices from Scn1aHet mice with a deficit in sodium channel NaV1.1, an important channel for interneuron excitability, have more seizure-like events (SLE) than wild-type (WT) littermates in a low Mg^2+^ model. Furthermore, we show novel data that demonstrate an increased seizure-propagation rate in the Scn1aHet mice, likely due to the well-documented decreased firing rates of parvalbumin-positive interneurons in these mouse models (Favero et al., 2018, Tai et al., 2014, Martin et al., 2010). We provide this new python-based software tool as an open-source, customizable solution for analysis and tracking of LFP activity using the 3Brain MEA recording system, but it can easily be adapted to any MEA recording platform. This GUI will also likely be suitable for analysis of large-scale EEG recordings and provide a useful mapping tool for in vivo LFP activity. Our current GUI has a particular utility for analysis of seizure-like activity but can be used for analysis of many other network LFP signals.

## 2 Methods

### 2.1 Ethical approval

All animal handling and experimentation involving animals were conducted following approved protocols according to the guidelines of the Canadian Council on Animal Welfare.

### 2.2 Slice preparation

Heterozygous *scn1a* knockout (Scn1a(+/-)) mice (Mistry et al., 2014) and WT littermates were used in this study. Heterozygous mice on the 129S6/SvEvTac background (MMRC strain number 037107) are crossed with C57BL/6 mice at The Jackson Laboratory. The pregnant mice are then shipped to Xenon Pharmaceuticals to litter. Pups are then genotyped to determine their genotype as either WT for the *Scn1a* gene (Scn1a+/+) or heterozygous for the *Scn1a* gene (Scn1a+/-). All mice used in the study were genotyped a second time on the day of euthanasia to reconfirm their genotype. Scn1a and WT mice were used in this study between the ages of P21-P28. Mice were housed in individually ventilated cages in 12 h light, 12 h dark lighting regime. Animals received food and water ad libitum. Mice were anesthetized with isoflurane before being euthanatized by cervical dislocation. Brains were then removed and stored in cold cutting solution (in mM): 3 MgCl2; 126 NaCl; 26 NaHCO3; 3.5 KCl; 1.26 NaH2PO4; 10 glucose. For multielectrode array recordings, 350 μm horizontal hippocampal sections were made, using a Leica VT1200 vibratome (Nussloch, Germany). Slices were then transferred to a holding chamber and incubated for 1–2 h at room temperature in artificial CSF (ACSF) containing (in mM): 2 CaCl2; 1 MgCl2; 126 NaCl; 26 NaHCO3; 3.5 KCl; 1.26 NaH2PO4; 10 glucose. All the solutions were bubbled continuously to saturate with carboxygen (95% O2 and 5% CO2).

Multi-electrode array recordings were performed on the 3Brain BioCAM DupleX system (Switzerland) using the 3Brain Accura HD-MEA chips with 4,096 electrodes at a pitch of 60μm. Slices were placed onto the electrodes with a harp placed on top to keep the slice pressed down gently to the recording electrodes. Slices were perfused continuously with artificial cerebrospinal fluid (ACSF) that had Mg^2+^ lowered to 25μM to induce epileptiform-like activity. Recordings were obtained from the entire slice, containing both the neocortex and the hippocampus. Experiments were performed at 33–36°C. The solutions were perfused at the rate of 5.0 mL/min. Signals were sampled at 10 kHz with a high-pass filter at 2 Hz.

### 2.3 Statistics

Statistics were done in GraphPad Prism 9.1.1 (San Diego, CA). Data was analyzed with unpaired Student’s t-tests, except for the analysis of seizure duration in Figure 7E, in which a Mann-Whitney test was used due to a significate F-test to compare variances. GraphPad Prism was also used to graph scatter-point data shown in Figure 7. Significance was set at P ≤ 0.05 for all analyses.

### 2.4 Data analysis and figures

The analysis platform and algorithms used were custom written in Python, including NumPy, pandas, SciPy, and visualizations using Plotly’s Dash libraries. The code and sample data files are provided through a GitHub repository^1^. Figures for the manuscript were created using Inkscape 1.1.

### 2.5 Local field potential (LFP) measures in channel groups

LFP count per second: To detect local field potential from voltage traces, the signal processing library from SciPy in Python is used, specifically the ‘scipy.signal.find_peak’ function. The inputs to this function include the ‘threshold’ and ‘width,’ which are received as inputs from the user in the GUI as Threshold (mV) and Time Duration (s) respectively. The ‘find_peak’ function returns the time index of peaks that exceed the minimum threshold value for the minimum specified duration for each individual channel; no maximum limits are set. The sum of count of the peaks for each channel in the group is calculated, which is then summed for all selected channels in the group to get the total LFP count for the group. This is divided by the time range (in seconds) of the signal to return the LFP count/s for the group. The LFP count/s for each channel is just the sum of the LFP count for individual channels divided by the time range. For channels in the measurement file or group of channels selected, active channels have more than 20 peaks in the selected time range. The channels are sorted in the decreasing order of LFP activity, and the top 20 most active channels are selected within the group of channels. The LFP count per second is also calculated for the 20 most active channels in the group in certain cases.

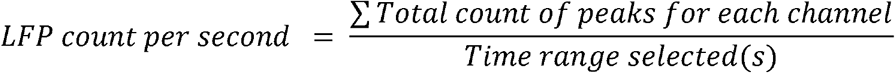

LFP mean amplitude (mV): For each channel, the absolute voltage amplitude at each peak is calculated, which is used to calculate the average LFP amplitude for each individual channel. For a group of channels, the LFP mean amplitude is the mean of average amplitude of peaks for individual channels. The LFP mean amplitude can also be calculated for the 20 most active channels in the group, rather than include all the channels in the group.

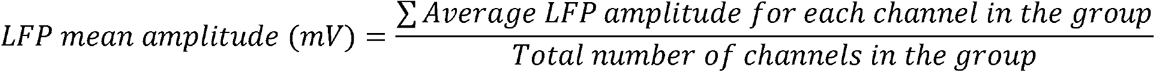

### LFP mean duration (s)

For each channel, the width of each peak is extracted using ‘properties’ in the ‘find_peak’ function, which is used to calculate the average LFP duration (width) for each individual channel in seconds. For a group of channels, the LFP mean duration is the mean of average duration of peaks for individual channels. The LFP mean duration can also be calculated for the 20 most active channels in the group, rather than include all the channels in the group.

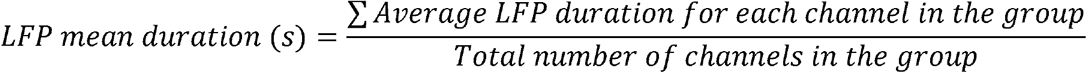

## 2.6 Seizure-like event (SLE) network measures

Maximum distance of spread of SLE: The Euclidean distance from the electrode at which the initiation of SLE is observed in the slice to the furthest point from the initiation point. The row and column number are used as the x and y coordinates respectively. The Euclidean distance between the x, y co-ordinates have no unit. It is multiplied by the electrode spacing in micrometer to get the distance of spread of seizure-like activity in the slice.

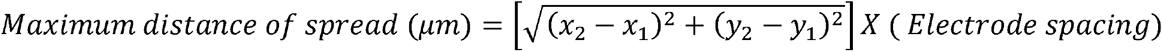

### Duration of SLE

This is calculated for each channel in a selected group. The difference between the end time and the start time of the seizure-like event in the selected time window of the ‘Channel Raster (Groups)’ gives the seizure duration for that channel. The mean and maximum duration are calculated for each group from the duration of seizure-like activity of all channels in that group.

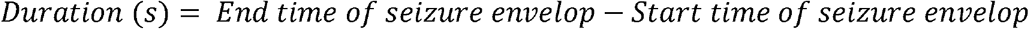

### Seizure rate

For the selected time interval in the ‘Channel Raster (Groups),’ the start time, end time of SLE are calculated for all channels in the group. The maximum distance of spread of the SLE is also calculated for that group. The seizure rate is the maximum distance of spread of the SLE divided by the mean difference in the start times of the seizure for each individual seizure.

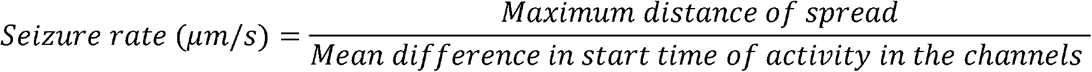

## 3 Results

The data processing pipeline for LFP activity and seizure analysis consists of three steps starting from the measurement file as shown in Figure 1. A typical measurement file consists of 4096 channels recorded for about 50 minutes at a sampling frequency of 10 kHz, in the hdf5 format, the file size is about 250 GB uncompressed. As a first step, channels that overlay the slice are selected based on the desired resolution, and exported using the 3Brain proprietary BrainWave4 software. This exported file consists of about 300 to 600 channels with the original sampling frequency, and a reduced file size of 80 GB. The file size and number of channels selected in this step can vary depending on the recording sampling frequency, resolution required for the analysis, and recording time. Second, the extracted channels that overlay the slice from the previous step are down-sampled in Python, this down-sampled file maintains the same data structure and hdf5 format as the original recording, thus has backward compatibility with BrainWave4 software. The down-sampled file is now ready for use with our custom interactive MEA Viewer - Xenon LFP Analysis Platform. The GUI is built in Python using the Plotly’s Dash library, which renders visualizations in a user-friendly web interface. A snapshot of the opening page of the web interface is shown in Figure 2. The analysis platform has the following key functions: 1) MEA Viewer Functions: This includes options to select and view individual channels, generate raster plots for all the channels, apply digital signal processing tools including FFT, low-pass, high-pass, and band-pass filters. 2) Channels Group Functions: This has options to select three different regions or groups of channels, apply peak detection, generate custom raster plots, apply digital signal processing tools, and generate summary measures including LFP count per second, number of active channels, mean LFP amplitude, and mean LFP duration. 3) Seizure Detection and Analysis Functions: This is an unsupervised automatic SLE detection on selected channel groups and analysis of metrics on seizures observed in the slice. Moreover, Python and Plotly’s Dash, which is based on object-oriented programming and reactive callbacks, provide options to customize, change the layout of visualization, analysis methods, summary measures, and data processing algorithms in the GUI, as per the user requirement within each of these functions. The GUI application is either hosted and run on a server or run in the local machine. While running the Python script in the local machine by default the application can be accessed using http://127.0.0.1:8050/ on a standard web browser.

**Figure 1:**
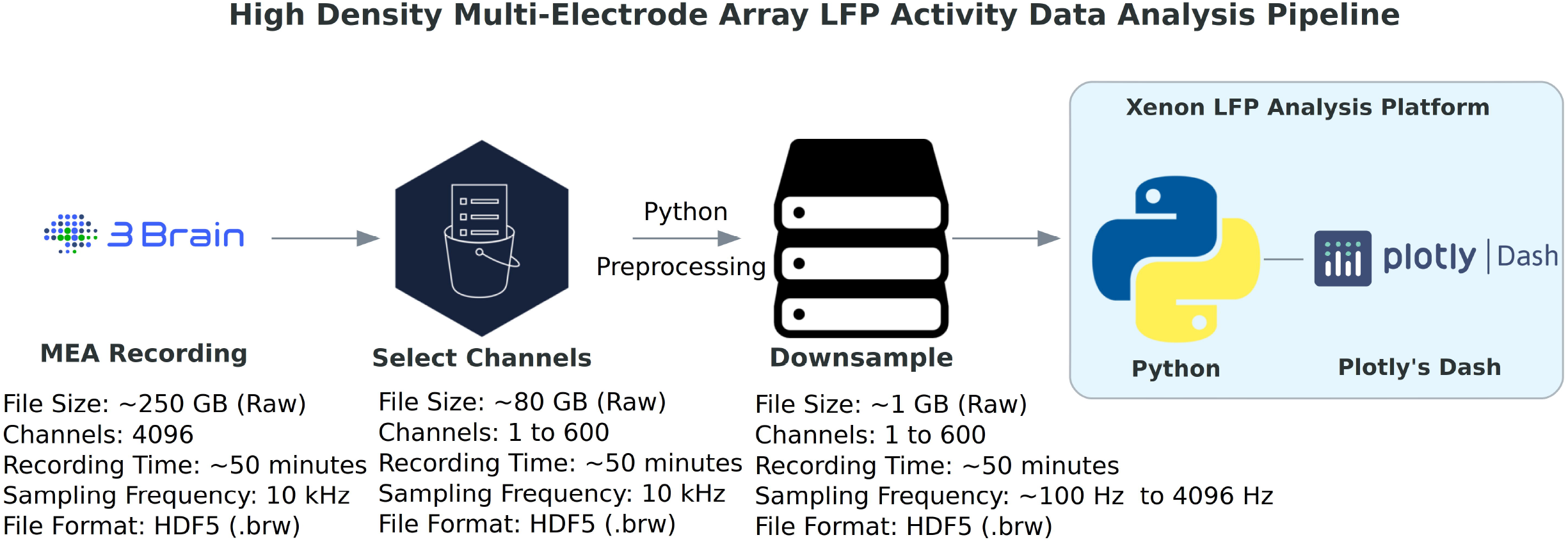
High-density multi-electrode array (HD MEA) data-analysis pipeline. The data processing for LFP activity detection and network analysis starts by selecting a group of about 600 channels that overlay the slice or region of interest, which are exported from the original hdf5 measurement raw file (4096 channels, sampled at 10 kHz) to a reduced hdf5 file. This reduced file is further down sampled from 10 kHz to a desired frequency. This will be the working hdf5 file for the Xenon LFP Analysis Platform. [3Brain Logo: © Copyright 3Brain AG, Python Logo: © Copyright Python Software Foundation, Plotly’s Dash Logo: © Copyright Plotly]

**Figure 2:**
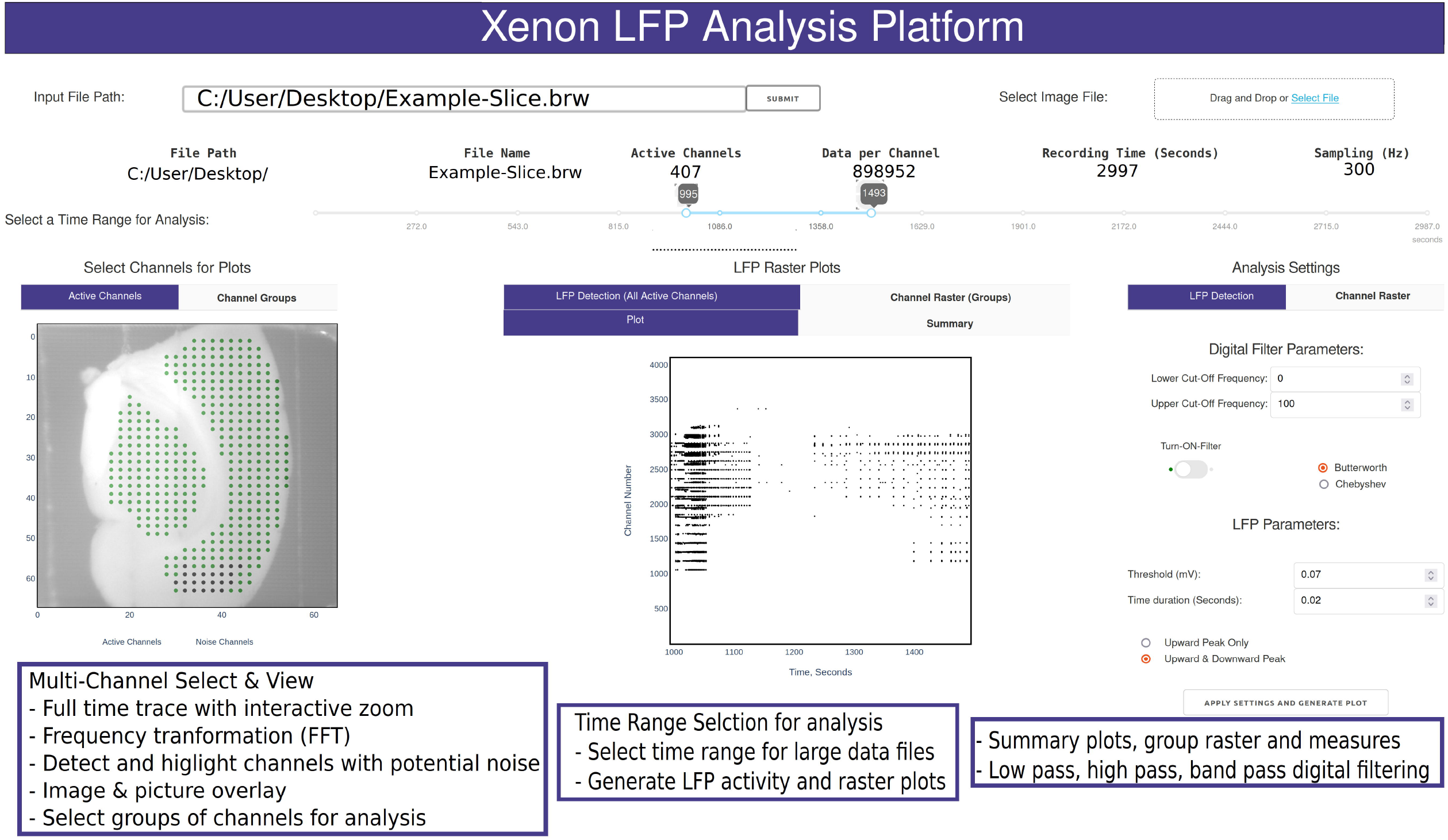
Snapshot of the analysis GUI features. A view of the analysis GUI which is rendered in a html browser built on Python using Plotly’s Dash. The GUI has several interactive features from individual and group channel selection, low-pass, high-pass, and band-pass filtering, viewing entire trace or a small section of the trace, Fast Fourier Transformation (FFT) of sections from selected traces, customized raster plots, small groups of channels, and generation of group summary measures.

### 3.1 MEA viewer functions

A common challenge among researchers using large MEA recording platforms is that it is not easy to explore the raw data. The Xenon LFP Analysis Platform functions are aimed to facilitate exploring the raw data, including viewing entire time-series, apply threshold detection, signal processing tools to individual and groups of channels. The entire platform is built for interactive explorations and analysis, while rendering the visualizations quickly in a few seconds. The time range selection (in Figure 2) is used to load and perform analysis on desired sections of the trace or the entire recording. Selecting channels for analysis is as easy as clicking using the mouse on the green dots that are shown with the slice image in the background (Figure 2). Each point corresponds to the channel location, x and y axis referenced to the original row and column number on the MEA array. Multiple channels can be selected by holding down the shift key while clicking using the mouse. The selected channels automatically load and display for the given time range. Any changes in channel selection, time range selection, or analysis settings dynamically change the analysis measures and output displays. The analysis setting can be used to apply digital low-pass, high-pass, band-pass filters, modify default threshold and duration for peak detection and raster plot generation. All plots are interactive; they can be zoomed in, zoomed out, downloaded as *.png files. Zooming small sections of the time-series in the LFP activity view will automatically generate FFT traces in an adjacent window. These functions are demonstrated in the supplementary video 1 (S1)^2^.

Figure 3 shows a sample analysis demonstrating the MEA viewer functions in detail. In this example, 407 channels are exported from the original recording for analysis. The analysis file was down-sampled from 10000 Hz to 300 Hz. The green dots overlay the neocortex, and the electrodes corresponding to the brown dots overlay the hippocampus (Figure 3A). The raster plot in Figure 3B highlights LFP activity in the entire recording. The channels are arranged according to their x, y position in the row and columns from 1 to 4096. The default threshold and duration for LFP activity is 0.07 mV and 0.02 seconds, however the raster can be regenerated for a range of input values by modifying the parameters in the analysis settings (see Figure 2 and S1), including generating raster after application of low-pass, band-pass, and high-pass filters. Figure 3C shows time-series traces from three electrodes (highlighted in Figure 3A); one from the hippocampus displayed in blue, and one from either end of the neocortex displayed in red and aqua, respectively. It is interesting to note the difference in the activity pattern in the three traces at the same instant of time. While Figure 3C show traces for duration of the recording from the selected electrodes, a section of these traces can be selected to view on a faster timescale (Figure 3C, AA; Figure 3D), as shown in Figure 3D. The black vertical markers at the top of each trace shows LFP activity detected based on the given threshold and duration. This further highlights the difference in the activity pattern in the different regions of the slice at the same time instance. We can apply digital filters to the traces, for example a 40-150 Hz band-pass filter to view low and high-gamma activity (Figure 3E). We see the blue and red trace have some gamma components, however the aqua trace does not have significant gamma components in the LFP activity. The time traces are interactive; to view spectrum plots (FFT), a small selection of the trace can be selected which automatically generates the FFT traces adjacent to the time-series traces (as shown in Figure 3E and Figure 3F). The filtered and original traces are usually overlaid, however, to view one or the other, clicking on the legend selects/deselects the trace to view one or both at a time. When digital filters are applied, the amplitude spectrum of the band-pass-filtered and unfiltered (purple) traces are overlayed to show the effect of filtering (Figure 3F, unfiltered: purple, filtered: electrode-specific colors).

**Figure 3:**
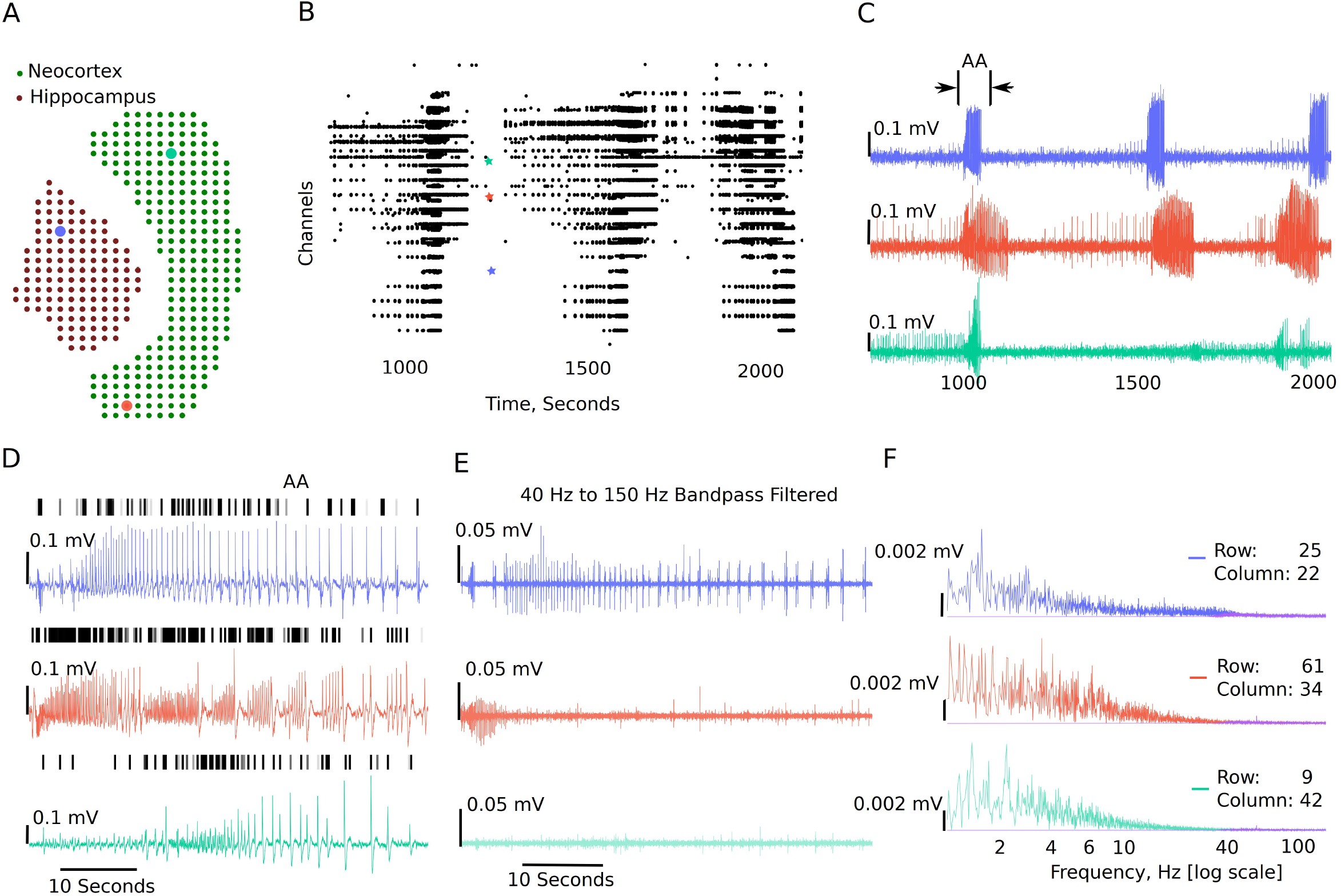
Example visualizations generated from the GUI including raster plots, time-series traces, LFP activity peaks, and time frequency transformations. **(A)** 407 channels selected for this analysis representing the MEA sensor spatial array, covering both the neocortex and the hippocampal regions. **(B)** Raster plot for all the channels in the working file irrespective of brain regions for a selected time range, demonstrating time points of when the activity occurs in the slice. **(C)** Three selected traces, the blue trace being from the hippocampal region, the red trace being from one end of the neocortex, and the aqua trace being from the other end of the neocortex. **(D)** Zoomed in view of the first seizure (the region bracketed in **(C)** as AA), with the black dashes showing a peak find function within the GUI. **(E)** This demonstrates the ability to plot filtered traces along with the raw traces. **(F)** The amplitude frequency transformations for the traces in **(D)** and **(E)** (band-pass filtered), the filtered FFT spectrum for each is shown in purple.

### 3.2 Channel group functions

The channel group functions are aimed at comparing two or three different regions of the slice, and compare LFP activity summary measures, while also generating a raster plot to study difference in activity pattern in different regions. The analysis starts with the ‘Channels Groups’ tab (see Figure 2 and Supplementary video 2 (S2)^3^). Channels groups can be selected by clicking on channels or by using the box or draw tool to select multiple channels at the same time. The groups tab enables selecting channels under three groups (Group1, Group 2, and Group3). The channels for each group are selected under their respective tab. Once respective groups and channels are selected, analysis settings can be modified from the default followed by clicking on ‘Apply Setting and Generate Plots,’ which generates the raster plots and summary measures (Figure 4A and Figure 4B). The measures automatically calculated are also shown in Figure 4B, which include Total LFP count/s, total channels. Channels that have more than 20 LFP activity peaks count in the selected time interval are considered as active channels, and the last three measures are the LFP Count/second, mean amplitude, and mean duration for the top 20 channels in each group. As shown in the summary table, for Group 1 in Figure 4B (bottom), which includes 132 channels in the hippocampus, 83 channels are active, of which only the top 20 are used to compare the mean amplitude and mean duration. When we look at the raster corresponding to the channels in red (hippocampus), there is quite a bit of variability between channels, in the activity count, start times for seizure-like activity. To avoid bias, we select the 20 most active channels from each group to calculate and compare mean measures between the groups. Further, the total activity, LFP amplitude, and duration are shown in the summary plot that includes all the channels in the group. The plots and summary measures can easily be regenerated for suitable selection of the time intervals by modifying the time range selection, and channels in each group. The front-end table displays a consolidated summary for all channels in a group, however summary data for each individual channel is automatically generated into an excel file and saved in a results folder for further analysis. The saved analysis file consists of LFP count, mean amplitude, duration, channel number, and group number for each individual channel. This data is overwritten each time the analysis settings change to avoid creating multiple log files. To create a copy, the file can be renamed, or the code can also be modified to save all log files continuously in the results folder.

**Figure 4:**
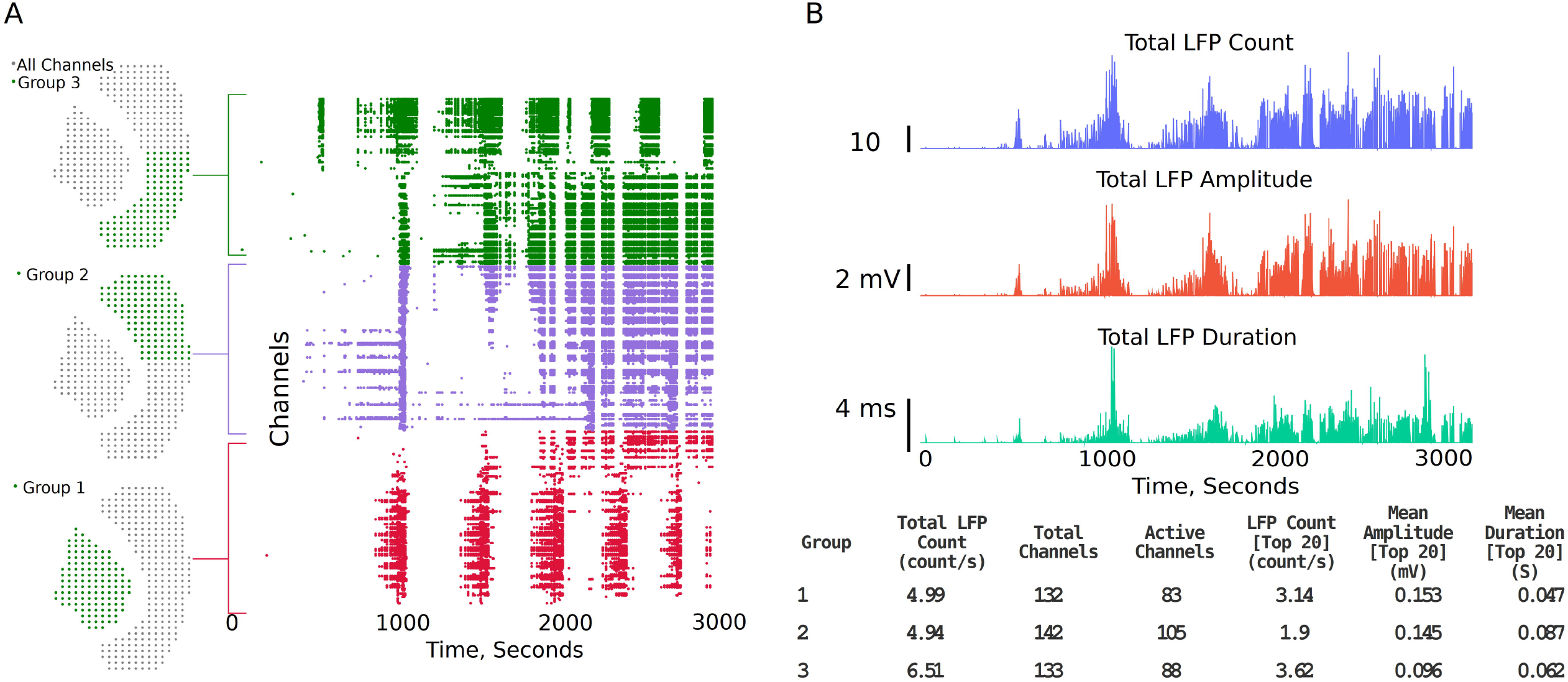
Channel groups and raster plot can be generated to visualize LFP activity in different regions of the slice. **(A)** Sensor locations corresponding to three different regions selected for analysis and the region-specific raster plots. Group 1 being the hippocampus, whilst group 2 and 3 each being one half of the Neocortex. **(B)** Summary plots and measures that can be generated within the analysis platform.

### 3.3 Seizure detection and analysis functions

#### 3.3.1 Seizure detection

Detection and classification of interictal, ictal or SLE can be quite challenging due to different types of epileptiform activity, variability from type of measurement paradigm (4-aminopyridine, low Mg^2+^, low Ca^2+^, high K^+^), and inherent experiment-to-experiment variability (Ghiasvand etal., 2020, Campos et al., 2018). In the Xenon LFP Analysis Platform, we introduce a simple method to detect SLE using changes in spectral activity and LFP activity in the traces. We found this method quick and easy to apply to many channels (>400 channels) at a time and compare different treatment effects. Moreover, this is efficiently implemented using numpy, scipy, and signal libraries in Python. The steps involved are illustrated in Figure 5.

**Figure 5:**
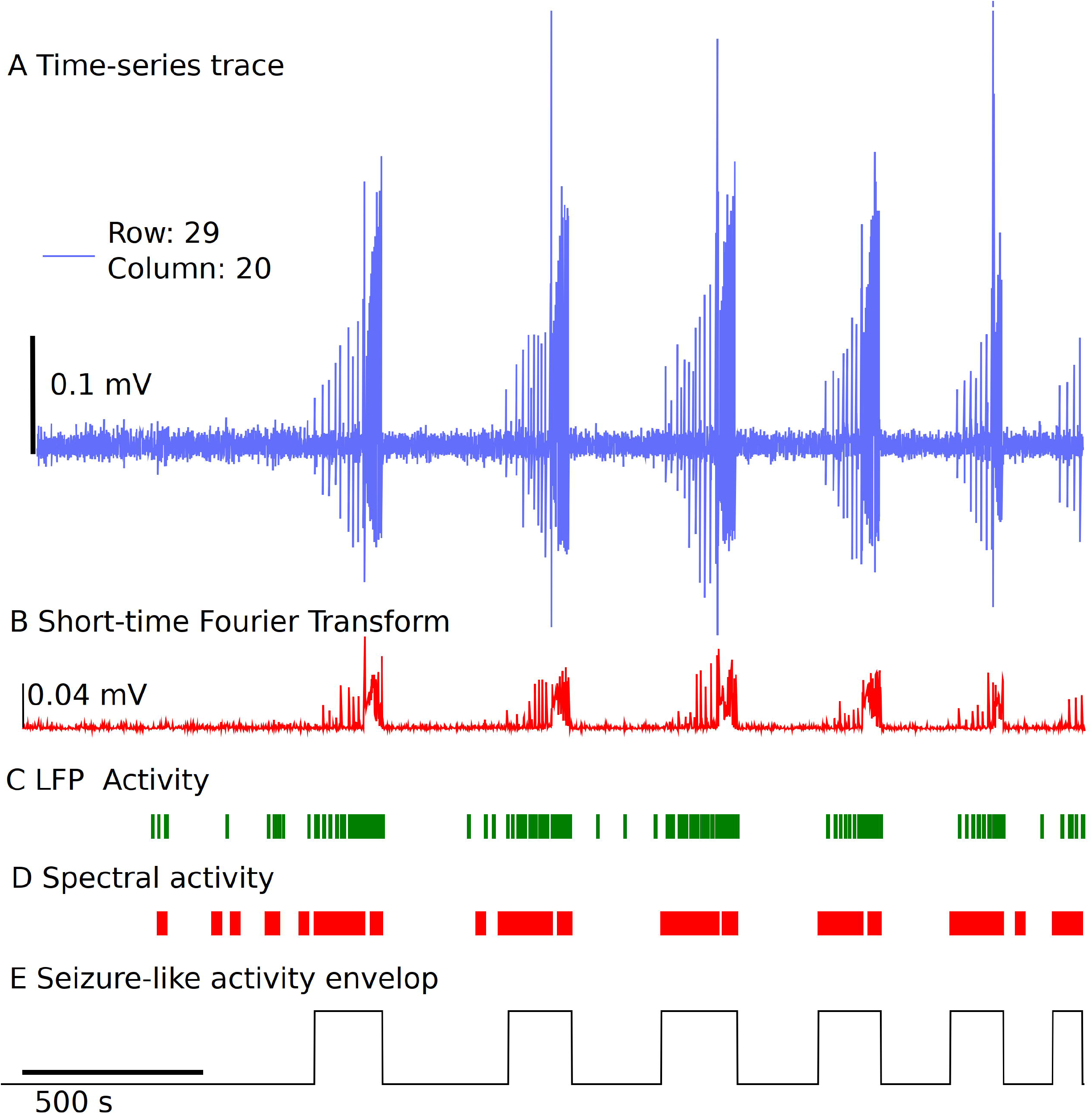
Simple and fast unsupervised seizure detection method. **(A)** The raw trace from the recording down sampled to 300 Hz frequency for a sample channel. **(B)** Spectral activity calculated from the Short-time Fourier Transform using Hanning Window, for a time window length of 1 second, with no overlap. **(C)** LFP activity detected using a threshold of 0.07 mV and 0.02 s, followed by applying two sets of sliding windows (length 30 datapoints, and 500 datapoints) to detect time regions of continuous activity. **(D)** Spectral activity detected when the magnitude is greater than 6 standard deviations from the reference spectrum. The spectral activity is also passed through two sliding windows to detect regions of continuous spectral activity. **(E)** The overlapping regions of LFP activity and spectral activity of 10 seconds or more are used to identify the seizure envelop. The start of seizure is primarily identified using the time point when the spectral activity is greater than 6 standard deviations from the baseline.

We start with the time-series trace down-sampled to 300 Hz, the first 60 seconds of the trace with no noise, and LFP activity is selected as a reference section of the trace to get a baseline for spectral activity. The spectral magnitude is calculated using the Short-time Fourier Transform (STFT), with a few variable parameters that can be set or standardized in the analysis platform, including length of time segment, window, and overlap points, which is shown in Figure 5B. Two sliding windows of dimension 30 datapoints and 500 datapoints are applied to the spectral activity peaks and LFP activity peaks independently, to detect regions of continuous seizure-like activity and time regions of no activity (Figure 5C and Figure 5D). This has a few parameters that can be standardized based on the experiment paradigm. In the examples discussed we use 6 standard deviations from the baseline spectral magnitude to detect high spectral activity and 0.07 mV voltage threshold, 0.02 s duration for LFP activity. The sliding window length (30 × 1 and 500 X1) and cutoff threshold parameter for automatically detecting spectrally active time regions post windowing can also be standardized. Once we have the time points of continuous spectral activity and LFP activity, we use overlapping points of both spectral activity and LFP activity to detect the seizure envelop. In general, the start of SLE is primarily detected when the spectral activity exceeds 6 standard deviations from the baseline and has continuous spectral and LFP activity for a minimum of 10 seconds. This again can be modified based on the experiment paradigm. If some seizures are closely spaced, parameters can be changed to a different value based on user preference. Once we have the seizure envelop with start and end times, we use this to further calculate the rate of seizure spread, distance of spread of seizure within a region of a slice using the group selection as discussed in the next section. This being an unsupervised method, and the variability of the nature of seizure-like activity in different regions of the tissue and between experiments, this may require manual verification by selecting a few channels and checking if the automatic envelop detect has good accuracy. We noticed that when the signal to noise ratio is high, and when clear LFP activity and spectral activity is detected, the algorithm performs well, but may need some adjustments to the parameters when the signal to noise ratio is low or LFP activity is not clearly differentiable.

#### 3.3.2 Seizure analysis

The channel group raster is required to perform the seizure detection and analysis. Each group has a separate tab (Supplementary video 3 (S3)^4^) under which individual channels can be selected to view seizure-like activity highlighted by the envelop (Figure 6A). Figure 6B demonstrates the raster plot for three different groups. Using the raster, a region can be selected with a potential SLE, as shown in Figure 6B (non-grey section), to generate summary measures and a visual of the channels that have a SLE within the selected section (Figure 6C). The channel dots highlighted in red are channels in the respective group that have a SLE, the blue dots are channels that did not participate in the SLE, while the grey dots have not been selected. The time interval shown in the summary table in Figure 6C is the selected time interval in the raster plot (Figure 6B -non-grey section). The distance, duration, and seizure rate are calculated from the start and end times of seizure envelop in each of the channels in the group for the selected (zoomed in) seizure.

**Figure 6:**
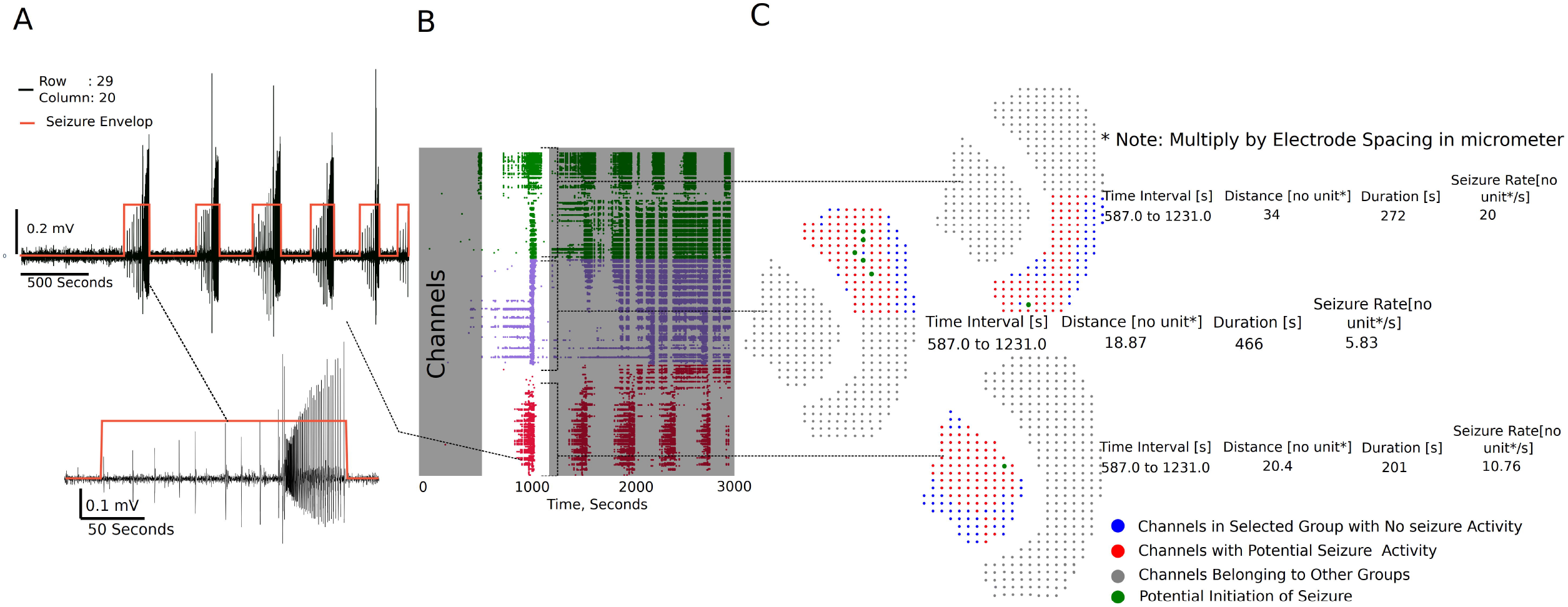
Seizure activity tracking over space and time. **(A)** Seizures in individual channels in a group are automatically detected, and their respective start and end times can be tracked across channels in that group. **(B)** Regions of the raster between time intervals can be selected as demonstrated to generate seizure maps of selected brain regions within the interval. **(C)** Seizure map for the time interval selected and channels in the group, including initiation sight of the seizure, maximum distance the seizure spread from the initiation point, duration of the seizure, and the rate of seizure spread across the tissue.

**Figure 7:**
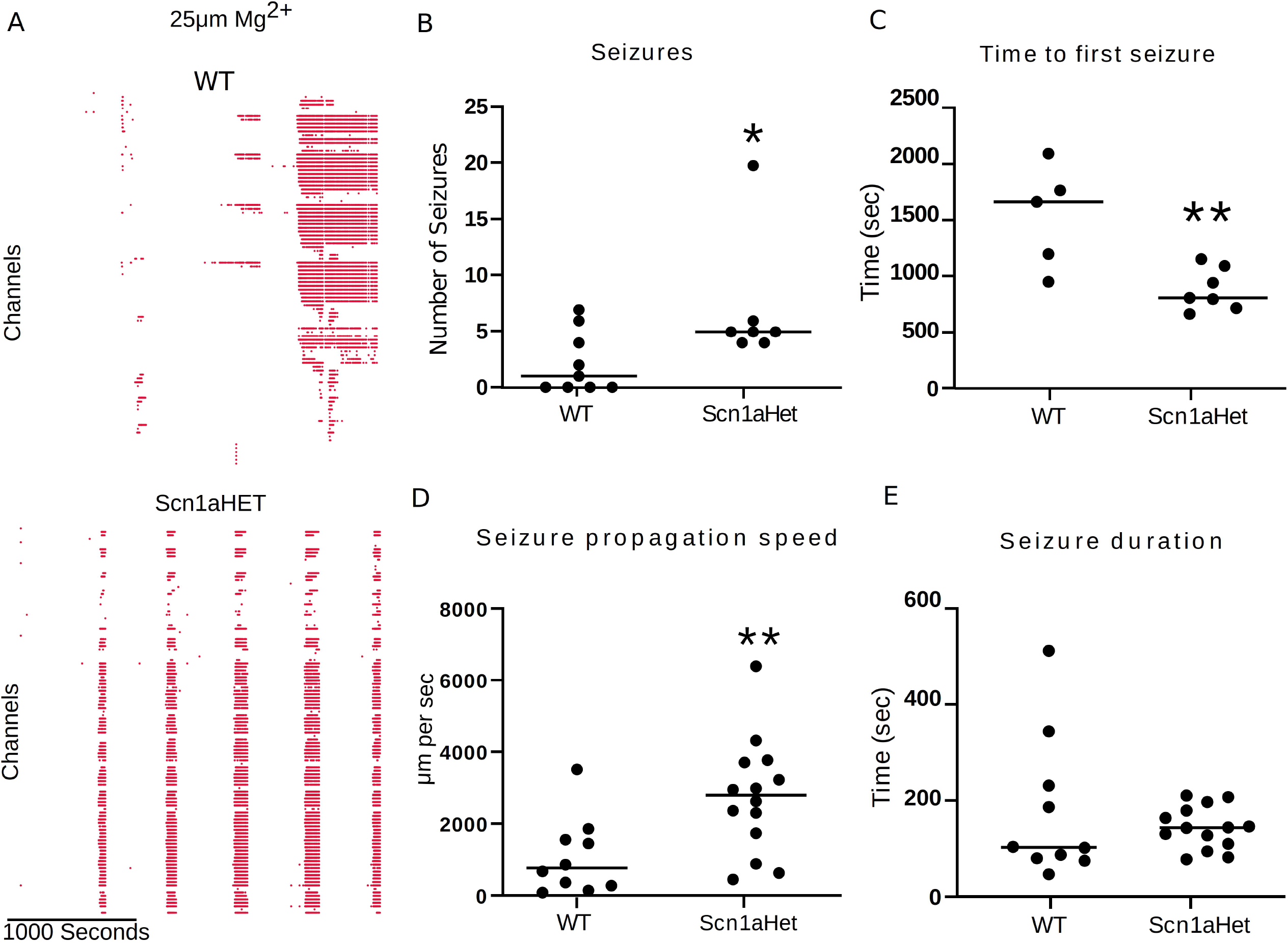
Scn1aHet mice have an altered seizure pattern in the low Mg^2+^ model. **(A)** Example raster plots from control slices and slices from Scn1aHet mice. **(B)** Scn1aHet have significantly more SLE than littermate controls (unpaired t-test, p = 0.04, n = 7-9). **(C)** From the slices that displayed SLE, Scn1aHet demonstrated a significate increase in time to first seizure compared to littermate controls (unpaired t-test, p = 0.006, n = 5-7). **(D)** SLE from the Scn1aHet propagate significantly faster than seizures in the littermate controls (unpaired t-test, p = 0.008, n = 10-14). The first two seizure from the slices that had SLE were used in this analysis. **(E)** Seizure duration was not different between Scn1aHet and littermate controls (Mann-Whitney test, p = 0.66, n = 10-14).

Three metrics are calculated from the seizure envelop for all channels in the group: distance of spread, duration, and seizure propagation speed. The spatiotemporal origin of the seizure within a group is identified as the channel that first had spectral activity above the set threshold, and this timestamp and the location of the channel is used to further calculate the distance and rate of the seizure spread. For example, in Figure 6C, it is the maximum distance from the green dot to the furthest red dots. If more than one channel is highlighted green, then they have similar start times, and the maximum distance from each point is calculated to find the overall maximum distance. The blue dots do not have a seizure-like event and are not included in the calculation. The x, y position on a 64X64 grid places the channels at 1 unit dimension from each other. The array spacing in micrometer is multiplied by the distance and seizure rate to get the final measure in micrometer and micrometer/second respectively.

We next used this seizure tracking function to examine if slices from Scn1aHet mice, which are heterozygous for NaV1.1, have an altered phenotype in the low Mg^2+^ model of epilepsy. The example raster plots demonstrate a likely difference in number of seizure-like events between WT littermates and the Scn1aHet animals (Figure 7A). Further analysis revealed that the Scn1aHet mice do have significantly more seizures than the WT littermates over the course of the 50-minute recording (Figure 7B). Furthermore, we found that the start time to first seizure-like event was significantly sooner in the Scn1aHet animals compared to controls, further demonstrating an increased seizure phenotype in animals with a deficit in NaV1.1 expression (Figure 7C). Using our novel seizures tracking algorithm within our GUI, we compared the speed of seizure propagation in slices from control mice verses the Scn1aHet mice. Interestingly, this analysis demonstrated a significantly faster rate of seizure propagation in brain slices from the Scn1aHet mice compared to control (Figure 7D). There was no significant difference found in the duration of the seizures between the control and Scn1aHet mice (Figure 7E). This data demonstrates novel phenotypic feature of the Scn1aHet mice, a decreased time to the appearance of the first seizure-like event, and an increased rate of seizure spread through the tissue, likely due to deficits in feed-forward inhibition provided by the somatostatin and parvalbumin interneurons (Parrish et al., 2019, Cammarota et al., 2013, Trevelyan et al., 2007). These new analysis features provided by the Xenon LFP Analysis Platform provide new and exciting ways to understand phenotypic differences in transgenic animals, understand how pharmacology impacts neuronal network activity over space and time, and is customizable to fit any researcher’s needs.

## 4 Discussion

The Xenon LFP Analysis Platform aims to produce a lightweight interactive application with high-quality visualization rendered on a web browser, using open-source libraries (Python & Plotly’s Dash) that can be standardized to an individual’s research requirements. In the examples shown in the results section, we provide a snapshot of simple visualization and signal processing tools, however this can be expanded and customized to include additional features as per the users’ requirements by building simple data analysis models/functions and rendering them using callbacks in Plotly’s Dash. The data models with Xenon LFP Analysis Platform enable creating summary measures for comparisons, and visualizations on the browser, creating an interactive toolbox for viewing millions of datapoints at a time, to extract meaningful results and conclusions from the measurements. Furthermore, the application is scalable to larger datasets, with the ability to build functions that selectively read from small chunks of data from the hdf5 array rather than load the entire dataset into memory for rendering on the browser. However, it should be noted one of the drawbacks of hdf5 files to store HD-MEA data is that using single dimension large arrays to store data makes indexing and selectively reading channels very inefficient and difficult to parallelize (Dragly et al., 2018, Rossant, 2016b, Rossant, 2016a). Most HD-MEA measurement systems use the hdf5 files system to record/write data to disk, which has its advantages for portability of data, but limits data analysis pipelines to parallelize signal processing tasks on distributed systems or multicore processors and GPUs. Some cases require reading the entire array to memory for extracting a group of channels to apply a band-pass filter or Fast Fourier Transforms (FFT). The current working file size on the Analysis Platform is limited by local system RAM. Future work can extend the current platform to include a data pipeline to work with larger files of 250 GB or more exceeding the system memory, using parallel computing algorithms for signal processing and visualization tasks including filtering, FFT analysis, spike sorting on a larger scale, which is a developing research area for computational neuroscience that requires more exploration (Jonathan W. Pillow, 2019, Street, 2021).

The spatiotemporal resolution of HD-MEA recordings on tissue slices provides high-quality data, while also presenting big data challenges in visualization and analysis, including extracting meaningful reproduceable results. This can further be complicated when testing long-duration drug protocols to include multiple compounds at different concentrations resulting in terabytes of data that can become overwhelming to analyze and compare (Perkel, 2018). There is always a need for simple data pipelines and new analysis platforms that are open source, user friendly, scalable, and portable that can produce repeatable analysis results for ease of comparison between paradigms and datasets (Sejnowski et al., 2014, Mouček et al., 2014). Standardization of analysis tools to compare different drug protocols is key to make sense of terabytes of data collected using different compounds, concentrations, and drug-treatment effects (Sobolev et al., 2014). The LFP Analysis Platform enables this by setting up standard functions with customizable parameters to generate raster plots and LFP metrics, this includes unsupervised methods to detect seizure-like activity. There are three key groups of measures: 1) summary measures relating to all channels in each recording, 2) metrics relating to channel groups, and 3) seizure-like event measures tracked for selected regions in the raster plots for channel groups. In the first step, LFP activity raster and activity count for all active channels are summarized in ‘LFP Detection (All Active Channels)’ tab (Figure 2). This data is not saved and is just rendered on the browser for viewing, which may be useful to quickly review the recording. This is also linked to the selected time range and analysis settings, including threshold, duration, and digital filter parameters. In the second step, channel groups or select regions of the slice may be selected along with a specific time range to generate custom raster plots, along with metrics like the number of active channels, LFP count, mean LFP amplitude, mean LFP duration (‘Channel Raster (Groups)’). As shown in the results section, this is particularly useful to compare different regions of the slice, or different time regions within a recording for drug treatment effects. In addition to viewing, all measures for individual channels in each group are automatically saved as a csv file in the background for further analysis. In the third step, the raster generated in step two can be used to select specific time points of activity to view and analyze LFP and SLE activity (Figure 5). These measures track network activity based on seizure envelops, start times, and end times. This being an unsupervised method, and due to the variability in measurement for different slices and protocols, user intervention may be required in some cases to check the activity envelop, and careful selection of activity regions in the raster plot. It is our hope that making this analysis platform fully open source will allow others to add functions that enhance its utility for all and aid in addressing limitation of this current GUI, such as aspects of the analysis requiring some user intervention and finding work arounds to streamline analysis of even larger data sets.

With the advent of larger recording systems, allowing for up to 6 slices and over 1000 channels per slice during a single recording session, tools like this GUI are timely. These new systems will allow for immense screening of transgenic animals to elucidate aberrant network behavior (Mackenzie-Gray Scott et al., 2022) and large-scale drug screening of biological tissue. Furthermore, there is need to understand with epilepsy and other disorders how different brain regions interact with each other when challenged in media that induces increased network activity or when stimulated electrically or optogenetically (Cela and Sjostrom, 2020, Rafiq et al., 2003, Codadu et al., 2019b). While we now have the recording platforms to facilitate these research needs, we are still limited by analysis tools, and here we directly address some of these needs in our GUI and set important groundwork for further developments within this platform. We also perceive that this GUI will be useful in other large-scale electrophysiological recording systems where the researcher wants to understand interactions between LFP activity at different recording sites over space and time. For example, it would be particularly interesting to visualize multichannel human EEG recordings within the framework of this GUI, which could provide easy and efficient visualization of channel recruitment during various behavioral states with the current built-in features and custom additions.

Overall, the Xenon LFP Analysis Platform introduces a standard approach to analyze large HD-MEA recordings, using high-quality visualization rendered on a browser, simple algorithms, and metrics, with lot of customizable features and options for researchers. We demonstrate the utility of this new analysis platform with ex vivo data and demonstrate a novel finding in a low Mg^2+^ model of epilepsy from Scn1aHet animals. Slices from the Scn1aHet animals display an increased rate of seizure propagation compared to slices from WT littermates. Using hundreds of channels to map spreading activity such as seizures adds another important tool in the hands of neuroscientists and will complement low-resolution traditional imaging techniques, such as Ca^2+^ imaging and dye-based voltage imaging. We hope this GUI will serve as a tool for collaborative work between research labs to contribute add-ons and share results and findings.

## Data availability statement

The raw data supporting the conclusions of this article will be made available by the authors, without undue reservation.

## Acknowledgments

We would like to thank the entire Xenon family for their support of this work.

## Author contributions statement

RP conceived this work. AM, NC, and RP designed the computational methods. AM wrote the code and designed the visualizations. RP collected the data. AM and RP analyzed the data. AM and RP wrote the manuscript. AM, NC, and RP edited and approved the final draft.

## Footnotes

1 https://github.com/MicroBrew09/xenon-lfp-analysis

2 https://youtu.be/Xpg_W8hEmCw

3 https://youtu.be/8T5q2_mpQDg

4 https://youtu.be/8XhgcPpj3Ek

